# Design of synthetic activators for programmable gene expression

**DOI:** 10.64898/2026.10.19.760300

**Authors:** Wei Chen, Ruopu Jiao, Brian Coventry, Xiangli Qu, Magnus Bauer, Long Tran, Tabitha Tcheau, Kejia Wu, Zihao Song, Nicole Roullier, Xinting Li, Cameron Glasscock, Isaac B. Hilton, David Baker

## Abstract

Transcription factors control gene expression by recruiting transcriptional coactivators through intrinsically disordered transcriptional activation domains (TADs), whose weak and promiscuous interactions make it difficult to establish quantitative relationships between coactivator recruitment and transcriptional output, and to precisely modulate gene expression. To overcome this limitation, we set out to design coactivator-recruiting proteins using AlphaFold3-predicted TAD-coactivator interactions as a guide. We developed a high-throughput functional screening platform in mammalian cells and identified over 300 well folded nanomolar affinity binders to the transcription coactivators Mediator (multiple subunits), BRD4, NCOA, and p300; targeted interface mutations enabled systematic tuning of their binding affinities. Using a panel of p300 binders spanning nanomolar to micromolar affinities, we found an ultrasensitive relationship between coactivator-binding affinity and transcriptional output, with a sharp transcriptional activation threshold at ∼500 nM. Incorporating p300 binder designs into CRISPRa yielded compact synthetic effectors that drove stronger activation than VPR across multiple genomic loci with minimal detectable cytotoxicity. Combining p300- and Mediator-specific binders further enhanced transcription, revealing functional synergy between these coactivators. Together, our work establishes a quantitative framework linking coactivator-binding affinity to transcriptional output and provides compact synthetic effectors for programmable gene activation.

## Introduction

Transcription factors (TFs) regulate gene expression by binding DNA through their DNA-binding domains and recruiting transcriptional cofactors through effector domains. Although genomic studies have extensively mapped TF binding across the genome^1^ and high-throughput screens have identified hundreds of protein fragments capable of activating or repressing transcription^2–4^, how cofactors are recruited and how their combined activities shape the transcriptional output remains poorly understood. During gene activation, transcriptional activation domains (TADs) recruit coactivators from several major regulatory classes, including chromatin remodelers, histone modifiers, and the Mediator complex, which act at different steps to promote productive transcription (**Figure 1a**). TADs can be fused to DNA-binding domains or dCas9 to direct transcriptional activation, and combining multiple TADs, as in VPR, NFZ, and NMS, can substantially increase transcriptional activity^5–7^. This enhancement could arise from multivalent recruitment of the same coactivator, recruitment of distinct coactivators acting at different regulatory steps, or both, but their relative contributions remain unclear. Distinguishing these possibilities requires independent control over individual coactivator interactions. Native TADs are poorly suited for this purpose because they are intrinsically disordered and often engage multiple coactivators through weak interactions^8–10^, making the identity, affinity, and stoichiometry of recruited coactivators difficult to control. Consequently, it remains difficult to determine how the identity and strength of individual coactivator interactions, alone or in combination, shape transcriptional output.

**Figure 1.**
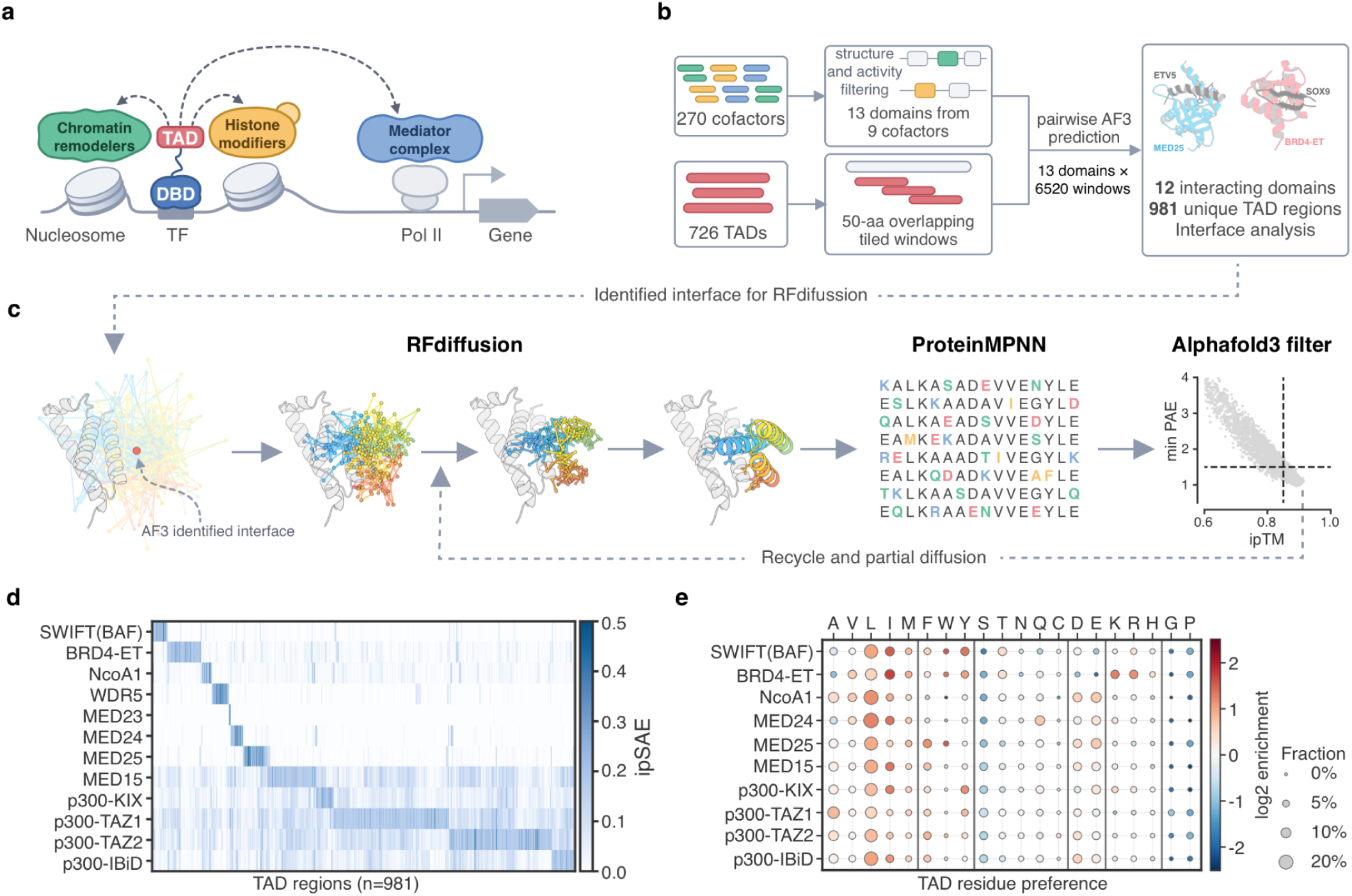
Structural mapping of TAD–coactivator interactions guides de novo protein design. **a.** Schematic of transcriptional activation. DNA-bound transcription factors (TFs) recruit transcriptional coactivators through transcriptional activation domains (TADs). TADs engage multiple classes of coactivators, including chromatin remodelers, histone modifiers, and Mediator, whose combined activities shape transcriptional output. **b.** AF3-based mapping of TAD–coactivator interactions. A set of 270 transcriptional coactivators was grouped by functional class and protein complex, and representative coactivators were tested for transcriptional activity. 13 domains from 9 coactivators were prioritized for AF3 prediction against 726 TADs identified from recent high-throughput screens. TADs were tiled into overlapping 50-aa windows to minimize sequence-length effects. This analysis identified 981 nonredundant TAD regions predicted to interact with at least one coactivator domain, whereas BRD4-BD1 showed no predicted interactions. Predicted complexes were further analyzed to identify interaction surfaces and sequence features for binder design. **c.** Schematic of the coactivator-binder design pipeline. AF3-predicted TAD-binding surfaces were used to define target interfaces for RFdiffusion-based protein generation. Designed structures were sequence-designed with ProteinMPNN and evaluated by AF3. High-confidence designs were further refined by partial diffusion, followed by a second round of sequence design and AF3 evaluation. **d.** Heatmap of the AF3-predicted TAD–coactivator interaction landscape across 981 TAD regions, scored using ipSAE. SWIFT, BRD4-ET, NCOA1, WDR5, and several Mediator subunits showed selective TAD-interaction profiles, whereas MED15 KIX and p300 domains displayed broader interaction preferences. **e.** TAD Amino-acid preferences at the predicted interfaces. Leucine and isoleucine were broadly enriched across coactivator-binding TADs, whereas individual coactivators showed additional sequence preferences. SWIFT- and BRD4-binding TADs were enriched for positively charged residues, and TADs interacting with other coactivators are enriched for negatively charged residues. Phenylalanine and tryptophan are enriched in MED25 interacting TADs and tyrosine enriched in p300-KIX-binding TADs.

We reasoned that replacing native TAD interactions with de novo designed proteins that engage individual coactivators through defined molecular interfaces could provide such control. Recent advances in deep-learning-based protein design, such as RFdiffusion^11,12^, enable the generation of structured, high-affinity binders against diverse target proteins. Here, we set out to develop a general strategy for designing coactivator-specific binders. As many transcriptional coactivators contain extensive disordered regions and lack complete experimentally determined structures, we aimed to first use structure prediction methods to characterize TAD-coactivator interfaces, and then to design binders targeting these interfaces (**Figure 1b, c**).

### Designing coactivator-specific transcriptional activators

To identify coactivator candidates for binder design, we grouped 270 transcriptional coactivators by molecular functions and transcriptional complexes, analyzed their structures, and tested representative coactivators for their ability to activate transcription (**Figure S1a**). Based on functional activity and the presence of exposed surfaces potentially accessible to TADs, we prioritized 13 domains from 9 coactivators for further analysis. We used AF3^13^ to systematically map predicted interactions between the 13 coactivator domains and 726 TADs recently identified through high-throughput screening^3,4,14^**(Supplementary Table 1)**. To accommodate variation in TAD length, we tiled each sequence into overlapping 50-amino-acid windows for AF3 prediction and identified consensus interaction regions as described in **Supplementary Note 1**. 12 of the 13 domains showed predicted interactions, yielding 981 nonredundant TAD interaction regions in total (**Figure 1b, Supplementary Table 1)**. Across the resulting interaction matrix, AF3 predicted differences in TAD-binding preference among coactivators. Many domains showed selective interaction profiles, including the SWIFT domain of the BAF complex, NCOA1, BRD4, and several Mediator subunits. In contrast, multiple p300 domains and the KIX domain of MED15 interacted more broadly across the TAD panel. However, even these more promiscuous coactivators showed nonuniform binding patterns, with clear preferences for subsets of TADs (**Figure 1d**). The predictions indicate that TADs engage different coactivators through short structured segments that adopt distinct conformations **(Figure S1c)**. β-strand conformations were frequently observed among SWIFT- and BRD4-binding TADs (81% and 71%, respectively), whereas interactions with the remaining coactivators were predominantly α-helical (89–100%) **(Figure S1c)**. The predicted α-helical interfaces shared common biochemical features: hydrophobic TAD residues contacted hydrophobic patches on the coactivator, while acidic residues frequently formed additional contacts with basic surface residues (**Figure S1d,e**). Beyond these shared features, individual coactivators showed differential enrichment for specific residues. For example, MED25 interfaces were enriched for phenylalanine and tryptophan (2.6- and 3.5-fold), whereas tyrosine was enriched ∼3-fold at p300 KIX but not MED15 KIX interfaces **(Figure 1e)**. The predictions also suggest that many domains can engage different TADs through distinct binding surfaces **(Figure S2 a-f)**. Together, these structural and sequence analyses defined coactivator binding surfaces and key interface residues.

We selected 10 coactivator domains (**Supplementary Table 2**), and used RFdiffusion to design coactivator binder proteins that target the interface residues identified in the TAD predictions above **(Figure S2 a-f)**. For each target interface, 3–4 residues recurrently contacted by different TADs were selected as design hotspots (for the BRD4-bromodomain AF3 did not predict interactions with any of the TADs examined, and hence hotspots were manually defined), and binder length was set to 60–100 amino acids. Each hotspot combination was used to generate 1,000 binder backbones with RFdiffusion^11,12^, followed by 10 sequences per backbone using ProteinMPNN^15^. The resulting binder–target complexes were evaluated with AF3 and filtered by minPAE and ipTM (minPAE < 1 or 1.5 and ipTM > 0.85) to prioritize high-confidence interfaces. Retained backbones were further refined by partial diffusion: limited noise was introduced and then removed by iterative denoising to optimize interface contacts, followed by another round of sequence design and AF3 evaluation **(Figure 1c)**. This workflow yielded approximately 3,000–5,000 filtered in silico designs per target, totaling ∼40,000 designs. These designs were synthesized and pooled for experimental screening.

### Functional screening identifies coactivator-specific activators

A major challenge in protein design is identifying functional candidates among designs that pass in silico filters. Conventional yeast-display screening presents binder libraries on the yeast surface and enriches target-binding variants by fluorescence-activated cell sorting (FACS), typically using purified recombinant targets. However, binding under these conditions may not translate into function in the crowded intracellular environment, where endogenous proteins can compete for binding or serve as off-target interaction partners. We therefore developed a pooled mammalian screening platform that directly selects for the intended function of these binders: transcriptional activation (**Figure 2a**).

**Figure 2.**
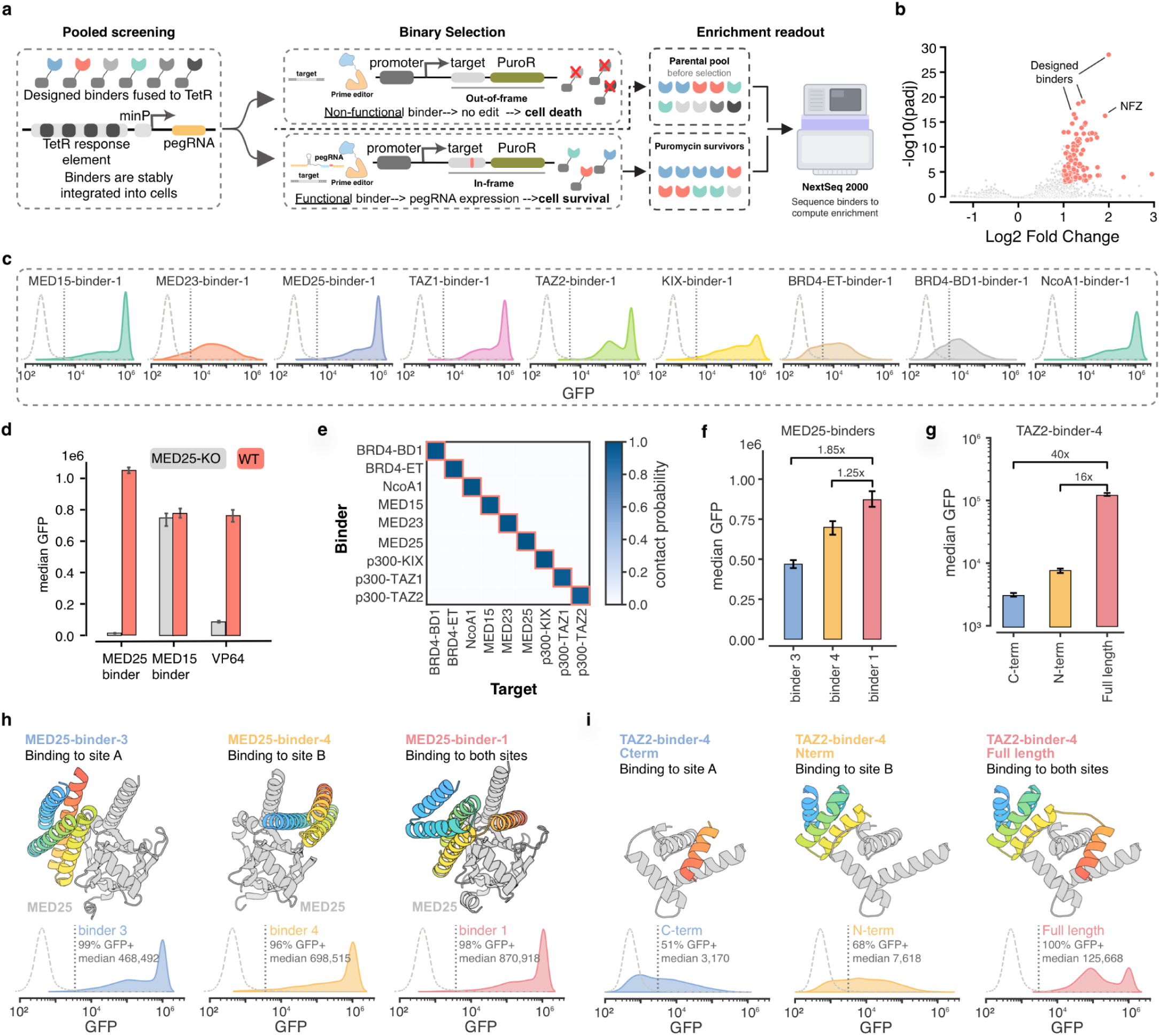
High throughput evaluation of designed binders. **a.** Overview of the molecular recorder-based screening platform. A binder library is fused to rTetR and integrated by PiggyBac transposition. With doxycycline, a functional binder activates pegRNA expression, recording its activity and restoring a downstream PuroR reading frame so only cells with functional binders survive selection. Abundance is quantified by sequencing genomic DNA from pre- and post-selection cells. **b.** Volcano plot of differentially recorded activity of designed binders. Red points indicate significant and substantial enrichment (Wald-test with Benjamini-Hochberg correction P <0.001 for fold-difference >2). Designed binders were successfully enriched together with positive control NFZ. **c.** Histogram of GFP fluorescence intensity for the indicated binder activators. Binders were fused to the C terminus of a zinc-finger DNA-binding domain and co-transfected with a GFP reporter containing the corresponding zinc-finger binding site. GFP fluorescence was quantified by flow cytometry. **d.** Specificity of the designed MED25 binder. Reporter and binder-expression plasmids were co-transfected into wild-type and MED25-knockout (KO) cells. A MED15 binder and VP64 were included as controls. The MED25 binder strongly activated reporter expression in wild-type cells but showed no detectable activity in MED25-KO cells. **e.** AF3 predicted specificity of all binders. Binder–coactivator pairs with predicted contact probability <0.5 were masked. **f.** Quantification of reporter activation by MED25 binders targeting interface A, interface B, or both interfaces. Data are quantified from the flow-cytometry distributions shown in **h**. **g.** Quantification of reporter activation by the full-length TAZ2 binder and its N- and C-terminal fragments. Data are quantified from the flow-cytometry distributions shown in **i**. **h.** Predicted binding modes of MED25 binders engaging interface A, interface B, or both interfaces (top), with corresponding GFP fluorescence distributions from the reporter assay (bottom). Binders are shown in color and MED25 is shown in gray. **i**. Predicted structure of the dual-interface TAZ2 binder and its N- and C-terminal interaction regions (top), with corresponding GFP fluorescence distributions for the full-length binder and separated fragments (bottom). Binders are shown in color and Taz2 is shown in grey.

For pooled screening, we adapted ENGRAM^16^, a prime-editing-based molecular recorder that couples transcription-dependent production of prime editing guide RNAs (pegRNAs) to defined insertions in genomic DNA. We coupled this recording mechanism to a frameshifted puromycin-resistance cassette, such that a 5-bp insertion into the genomic DNA tape restores the functional reading frame (**Figure 2a**). Transcriptional activation is thereby converted into a heritable genomic edit that confers puromycin resistance, enabling survival-based selection followed by sequencing-based quantification of library enrichment. Compared to other reporter-based assays, this recorder assay doesn’t rely on the expression level of the marker gene to enrich for the designed variants, and as such even weak gene activation can be captured.

We first engineered HEK293T recorder cells containing a Tetracycline Response Element (TRE) ENGRAM cassette, a prime editor expression cassette, and a genomic DNA tape with a frameshifted puromycin-resistance ORF. Candidate binder sequences were cloned as C-terminal fusions to reverse TetR (rTetR), enabling doxycycline-dependent recruitment to the TRE to control recorder activation. The pooled library, together with NFZ as an internal positive control, was integrated into recorder cells by piggyBac in triplicate. Recording was initiated seven days after transfection to allow dilution of residual episomal plasmids. Cells were cultured with 1 µg/mL doxycycline for two days, followed by selection with 4 µg/mL puromycin for five days. Cells were harvested before and after puromycin selection, and genomic DNA was extracted for sequencing the abundance of the integrated binders. Sequencing counts were analyzed using DESeq2 to identify significantly enriched designs. Candidate binders were enriched alongside the positive controls, with enrichment levels varying across designs (**Figure 2b**). Designs showing statistically significant enrichment of at least twofold (log2 fold change >1 and adjusted P <0.001) were selected for further characterization.

We performed four pooled screens covering 10 design campaigns against different coactivator targets (**Figure S3a-c**). Screening results were highly reproducible across independent integration replicates (Pearson’s r = 0.98, **Figure S3d-f)** and identified more than 300 candidate binders. The binders were predominantly α-helical, with BRD4-ET binders forming β-strand-mediated interfaces **(Figure S3g)**. We also identified binders against BRD4-BD1, despite the absence of predicted TAD interactions **(Figure S3g)**. Compared with native TAD fragments, the designed binders formed more extensive interfaces with their target coactivators. They buried more surface area (median 2,298 Å² versus 1,720 Å²; 1.1–2.6-fold across targets) and formed ∼2-fold more hydrophobic contacts (median 16 versus 7 residue pairs; 1.3–4.3-fold across targets, **Figure S3h**). Polar interactions were also markedly enriched in the designed interfaces (medians of 5 hydrogen bonds and 4 salt bridges, **Figure S3h**), whereas native TAD interactions contained few hydrogen bonds and frequently lacked salt bridges. Together, these features provide a structural basis for enhanced affinity and potentially greater interaction specificity. We selected 34 binders targeting 10 coactivator domains for individual validation, and all tested hits activated reporter transcription, with varying activation strengths (**Figure 2c; Figure S4a**). Of note, MED23 and BRD4 binders showed moderate activation, consistent with the relatively low activity observed upon direct recruitment of the corresponding coactivators.

We next assessed target specificity using genetic and computational approaches. For MED25, we generated MED25-knockout (MED25-KO) cells and compared MED25 binders with a MED15 binder and VP64. All constructs induced strong transcriptional activation in wild-type cells. In MED25-KO cells, the MED25 binders showed no detectable activation, whereas the MED15 binder remained active (**Figure 2d**). VP64 retained moderate activity, consistent with its promiscuous interactions of multiple coactivators^8^. Together, these results demonstrate strong MED25-dependent transcription activation by MED25-binders. To extend the specificity analysis to other targets, including essential coactivators for which knockout validation was impractical, we used AF3 to predict binder interactions across the coactivator panel. Predicted contact probabilities were high for the intended targets and low for alternative coactivators, supporting preferential recognition of cognate targets within the tested panel (**Figure 2e, Figure S4b**).

Many transcriptional coactivators contain more than one TAD-interaction interface **(Figure S2)**. We therefore asked whether these interfaces function independently or can act combinatorially. Our designed binders revealed examples of both behaviors. MED25 contains two TAD-interaction interfaces^17^, and we obtained binders targeting either individual interface as well as a binder engaging both **(Figure 2h)**. Engagement of either interface alone was sufficient to drive strong transcriptional activation, whereas simultaneous engagement of both produced only a modest additional increase in activity, corresponding to 1.85- and 1.25-fold increases over engagement of site A or site B alone, respectively **(Figure 2f,h)**. In contrast, one TAZ2 binder provided a distinct example of cooperative engagement. This binder contacted TAZ2 through two spatially separated interfaces: an N-terminal helical region engaged one surface of TAZ2, while a C-terminal protruding helix contacted a second surface. To determine the contribution of each interaction, we divided the binder into N- and C-terminal fragments corresponding to the two interfaces and tested them individually and in combination. Both isolated interfaces supported only weak activation, whereas the full-length binder showed 16- and 40-fold greater activity than the N- and C-terminal fragments, respectively **(Figure 2g,i)**. These results suggest that simultaneous engagement of the two TAZ2 interfaces strongly enhances functional p300 recruitment, which can be explained by cooperative multivalent interactions.

### Designed binders reveal ultrasensitive transcriptional regulation

Transcriptional regulation often involves multiple TFs acting together at the same locus, with their combined inputs shaping transcriptional output. These joint effects can arise through TF–TF interactions that enhance DNA occupancy^18^, as well as through their combined effects on coactivator recruitment^19–21^. TFs can jointly recruit coactivators in different ways: they may engage the same coactivator interface, effectively increasing local binding-site concentration; engage distinct interfaces within the same coactivator or coactivator complex, enabling multivalent and potentially cooperative recruitment; or recruit different coactivators, whose combined activities can act additively or synergistically on transcription **(Figure 3a)**. Although cooperative coactivator recruitment has been proposed, the quantitative relationship between TF–coactivator binding affinity and transcriptional activity remains poorly defined. In particular, how changes in binding affinity are translated into transcriptional output, and where this relationship begins to saturate, have not been systematically quantified. Here, we combine biophysical modeling with direct measurements of binder affinity and transcriptional activity to quantitatively link coactivator binding to transcriptional activation **(Figure 3b)**.

**Figure 3.**
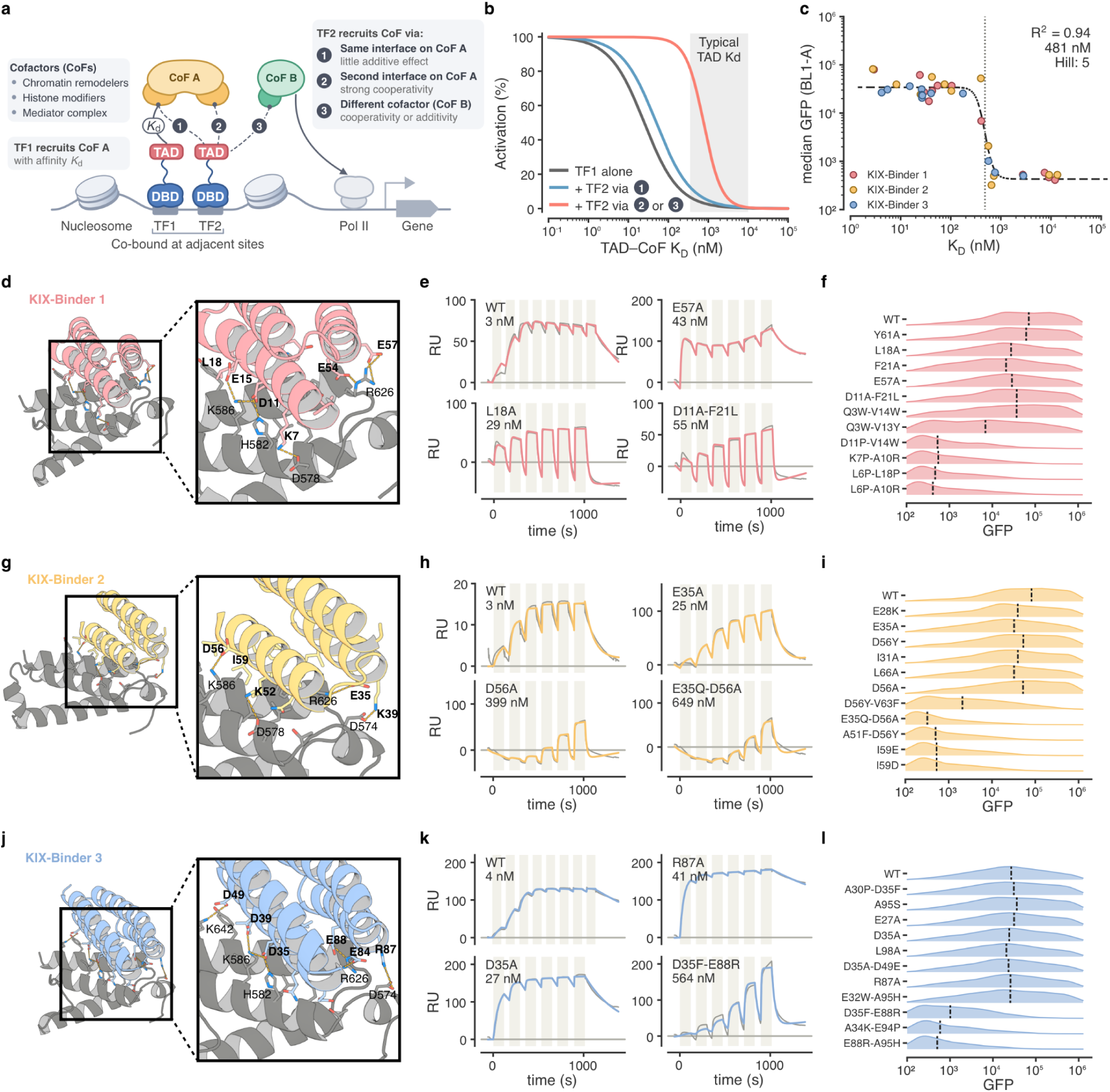
Designed binders reveal ultrasensitive responses in transcriptional regulation. a, Schematic of three modes by which TFs can jointly recruit transcriptional coactivators. TFs may engage the same interaction surface on a coactivator, increasing the local concentration of coactivator-binding motifs; engage distinct interfaces on the same coactivator or coactivator complex, enabling simultaneous multivalent binding; or recruit different coactivators that can act cooperatively within the same regulatory assembly or at distinct steps of transcription. **b,** Kinetic simulation of coactivator occupancy as a function of TF–coactivator binding affinity (K_D_) for a single TF (gray), two TFs engaging the same coactivator interface (blue), or two TFs engaging distinct interfaces on the same coactivator or coactivator complex (red). Coactivator occupancy was assumed to directly determine transcriptional activity. Shading indicates the typical affinity range of native TAD–coactivator interactions. Within this range, a single interaction or two TFs competing for the same interface produces limited occupancy, whereas simultaneous engagement of distinct interfaces enables efficient cooperative recruitment. **c.** Quantitative relationship between KIX-binding affinity and transcriptional activity for the affinity-tuned variants of KIX-Binders 1–3, using the SPR measurements and reporter activities shown in **d–l**. Median GFP fluorescence is plotted against the measured K_D_. The dashed line shows the fitted affinity–activity response, with a transition at 481 nM, an effective Hill coefficient of 5, and R^2^ = 0.94. **d–l**, Structural, biochemical, and transcriptional characterization of affinity-tuned KIX binders. **d, g, j,** Predicted structures of KIX-Binders 1–3 bound to the KIX domain, respectively. Enlarged views highlight key polar and hydrophobic interactions targeted for mutational perturbation. Binders are shown in color and KIX in gray. **e, h, k,** Representative SPR sensorgrams for the parental binders and selected interface variants of KIX-Binders 1–3, respectively. The measured K_D_ values are indicated for each variant. **f, i, l,** Histogram of GFP fluorescence intensities for cells expressing the parental binders and affinity-tuned variants of KIX-Binders 1–3, respectively. Dashed vertical lines indicate the median GFP fluorescence for each variant.

We began by modeling how TF–coactivator binding affinity influences coactivator occupancy across three recruitment architectures. A single TF recruits the coactivator efficiently only at high affinity, with substantial occupancy emerging in the single-digit nanomolar range. Adding a second TF engaging the same coactivator interface produces only a modest increase in recruitment, and transcriptional activity is again low at weaker binding affinities. In contrast, two TFs engaging distinct coactivator interfaces show strong cooperative recruitment, enabling efficient coactivator occupancy even at submicromolar affinities **(Figure 3b)**. Given that native TAD–coactivator interactions typically fall in the submicromolar-to-micromolar range^22–24^, these results suggest that cooperative recruitment could substantially enhance coactivator occupancy at physiologically relevant affinities, although additional mechanisms may also contribute in cells.

To test the predicted cooperativity in coactivator recruitment and quantify its strength, we took advantage of the well-defined interfaces of our designed binders to systematically perturb intermolecular contacts, tune binding affinity, and measure the resulting transcriptional activity. We considered both Mediator complex and p300 as candidate systems as they provide multiple TAD-interaction surfaces. However, recruitment through MED15 or MED25 alone already produced strong transcriptional activation in our assay, leaving limited dynamic range to resolve additional cooperative effects. p300 therefore provided a more suitable system, and we focused on the KIX domain because it was experimentally tractable, whereas TAZ1 and TAZ2 were difficult to purify and characterize biochemically. We therefore selected three KIX binders (KIX-binder-1–3) for systematic affinity tuning and functional characterization.

The three designs engage the KIX domain through extensive hydrophobic and polar contacts, including multiple hydrogen bonds and salt bridges that are less common in native TAD–KIX interactions **(Figure 3d,g,j)**. Salt bridges involving K586 and R626 were conserved across all three binders, suggesting that these contacts, together with hydrophobic packing in the KIX groove, contribute substantially to binding. We introduced targeted mutations to perturb either the polar interaction network or the hydrophobic interface, including disruption or charge reversal of key salt bridges and substitutions designed to disrupt hydrophobic packing. We generated 11 variants for each binder, yielding a panel of 36 proteins including the three parental binders. All proteins were expressed in *E. coli*, purified by size-exclusion chromatography, and were highly soluble and predominantly monomeric in solution (**Figure S5a**).

Binding affinities were measured by surface plasmon resonance (SPR) using biotinylated KIX immobilized on the sensor surface, with binder proteins injected at varying concentrations. The parental binders bound the KIX domain with K_D_ values of 3–4 nM, while TAZ1 binders failed to interact with the KIX domain **(Figure 3e,h,k, Figure S5b)**. Disrupting salt bridges with KIX residue R626 reduced affinity to 43 nM and 25 nM for Binder 1 E57A and Binder 2 E35A, respectively, while charge reversal in Binder 3 further weakened binding to 564 nM (D35F/E88R). Disrupting interactions with K586 reduced affinity to 55 nM, 399 nM, and 27 nM for Binders 1–3, respectively. The smaller effects observed for Binders 1 and 3 are consistent with additional acidic residues (E15 and D39, respectively) contributing to this electrostatic interaction network. Simultaneous disruption of both salt bridges in Binder 2 (E35Q-D56A) further weakened binding to 649 nM **(Figure 3h)**. Binder 3 also formed an additional salt bridge with D574, whose disruption reduced affinity to 41 nM **(Figure 3k)**. Replacing a core leucine with alanine reduced hydrophobic packing and lowered affinity to 29 nM for Binder 1 (L11A), whereas polar substitutions within the hydrophobic core significantly weakened binding to 8-14 μM (Binder 1 K7P-A10R and Binder 2 I59E/D, **Figure 3e, Figure S5 c-d)**. In total, we generated a panel of 36 KIX binders spanning more than three orders of magnitude in affinity, from low-nanomolar to micromolar (**Figure S5 c-e)**.

We fused the 36 variable affinity KIX binders to the zinc-finger DNA-binding domain and measured their transcriptional activity in the reporter assay. Transcription remained largely unchanged across affinities ranging from single-digit nanomolar to several hundred nanomolar, but dropped sharply as K_D_ approached the sub-micromolar range (**Figure 3c)**. Fitting binding affinity to transcription activity revealed a strongly nonlinear response, with an apparent transition around 500 nM and a Hill coefficient of 5. The affinity–activity relationship is ultrasensitive, with a steepness greater than expected from a simple two-site cooperative binding model **(Figure 3b)**, suggesting that additional cooperative or nonlinear steps after p300 recruitment may further shape transcriptional output. Together, these results support a model in which multiple weak TF–coactivator interactions can cooperate to achieve efficient coactivator recruitment and strong transcriptional activation.

### Designed coactivator binders enable compact and potent CRISPR activation

Given the high binding affinities and strong reporter activity of the designed binders, we next asked whether they could function as transcriptional activators at endogenous loci in a CRISPRa context (**Figure 4a**). Their compact size (65-85 amino acids) provides a practical advantage over conventional IDR-based activators such as VPR, which is approximately 600 amino acids (**Figure 4a**). We selected binders targeting several distinct transcriptional coactivators and fused them to the C terminus of dCas9 through flexible linkers. Plasmids encoding the dCas9–binder fusions were co-transfected with an sgRNA targeting the CD34 promoter, which is transcriptionally silent in HEK293T cells. 72 hours after transfection, CD34 surface expression was quantified by antibody staining and flow cytometry. All binders targeting p300 subdomains robustly activated endogenous CD34 expression (**Figure 4b**), with 11 of 12 binders generating a larger fraction of CD34-positive cells than VPR (**Figure 4c, Figure S6a, b**). The activated populations were also more homogeneous, with VPR showing 33% greater variability than the p300 binders, as measured by robust coefficient of variance (rCV) (**Figure S6c**). Despite this higher activation efficiency and uniformity, the magnitude of CD34 expression was generally lower than with VPR, with TAZ2-binder-4 as the only exception (**Figure 4b**). This distinction is consistent with the specific recruitment of p300 by the designed binders, whereas VPR can engage multiple classes of transcriptional coactivators, including Mediator and chromatin-remodeling machinery. Designs targeting NCOA1 or Mediator subunits MED15 and MED25 showed little activity at CD34 despite strong reporter activation, suggesting that recruitment of these coactivators alone is insufficient at a transcriptionally silent locus (**Figure 4b**)--in contrast, p300 recruitment can create a more transcriptionally permissive environment through acetylation of histones and other substrates^25,26^. To further validate that activation depends on p300 catalytic activity, we treated cells with A-485, a p300 catalytic inhibitor. A-485 abolished CD34 activation, confirming that binder-mediated activation requires p300 activity (**Figure S6 d,e**).

**Figure 4.**
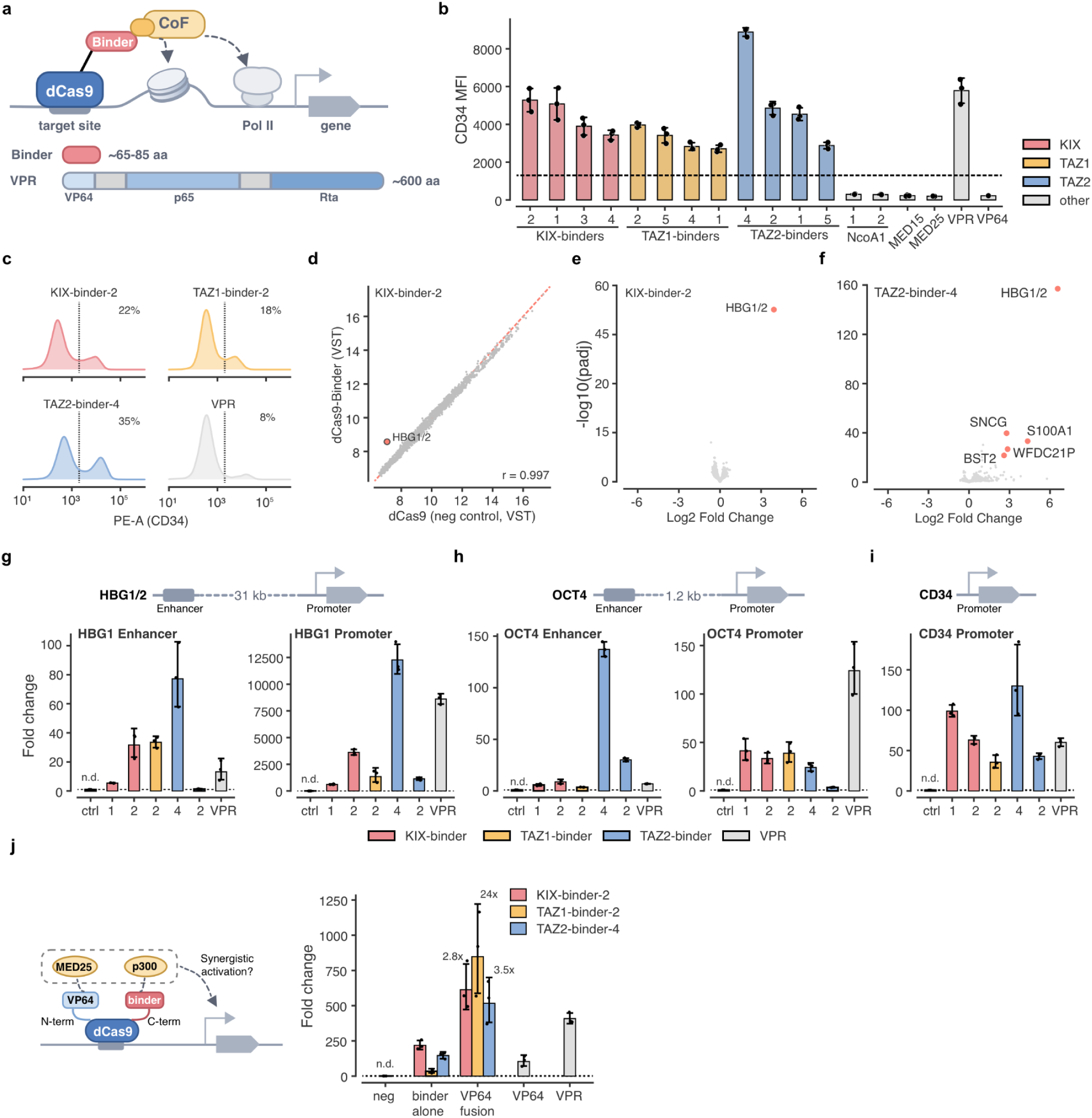
Designed binders function as compact and potent CRISPRa effectors. **a.** Schematic of dCas9-based recruitment of designed coactivator binders to endogenous genomic loci. The designed binders are 65–85 amino acids in length, substantially smaller than the tripartite activator VPR. **b.** Endogenous CD34 activation by dCas9-binder fusion. Binders targeting various coactivators were fused to the C-terminal of dCas9 and co-transfected into HEK293T cells with an sgRNA targeting the CD34 promoter. CD34 surface expression was measured by flow cytometry. KIX-, TAZ1-, and TAZ2-binders (p300 binders) robustly activated CD34, whereas NCOA1 and Mediator binders produced no detectable activation. **c.** Quantification of the fraction of CD34-positive cells for the indicated activators. Most p300 binders produced a larger activated cell fraction than VPR. **d,e.** Transcriptome-wide analysis of cells expressing the indicated dCas9–binder activators with an sgRNA targeting the HBG locus. Designed p300 binders strongly activated the targeted gene with limited transcriptome-wide perturbation. **f.** Volcano plot showing differentially expressed genes in cells expressing the TAZ2-binder-4 with sgRNA targeting the HBG locus. Red points indicate significantly differentially expressed genes (Wald-test with Benjamini-Hochberg correction P <0.001 for fold-difference >2). A small set of genes were also differentially expressed, including SNCG, S100A1, and WFDC21P. GO analysis didn’t reveal coherent functional enrichment of these genes. **g–i.** Endogenous activation across promoter and enhancer contexts. Selected p300 binders were fused to dCas9 and co-transfected into HEK293T cells with sgRNAs targeting promoters and enhancers of OCT4, HBG, and CD34 promoter. Target-gene expression was quantified by qPCR. Although activity varied across loci, p300 binders were generally comparable to or weaker than VPR when targeted to promoters, but consistently showed stronger activation when targeted to enhancers. **j.** Co-recruitment of p300 and additional transcriptional coactivators shows enhanced transcription activity. VP64 and p300 binders were fused to the N and C termini of dCas9, respectively. The dual activators were co-transfected into HEK293T cells with an sgRNA targeting the OCT4 promoter, and OCT4 transcript levels were quantified by qPCR. The dual-activator increased activation by 2.8-to 24-fold relative to the corresponding p300 binders alone and exceeded VPR-mediated activation at the OCT4 promoter.

To evaluate the target specificity of the designed activators, we performed RNA-seq in HEK293T cells expressing each binder together with a single sgRNA targeting the hemoglobin promoter. Across the binder panel, we detected strong transcriptional activation of the targeted gene (**Figure 4d, e, Figure S7a, b**). Transcriptomic profiles separated the activators into three groups: KIX- and TAZ1-targeting binders clustered together, TAZ2 binders formed a distinct group, and VPR showed a separate profile (**Figure S7c**). Overall, the designed binders produced moderately greater transcriptome-wide perturbation than VPR, although few genes were differentially expressed (≥2-fold change and adjusted P < 0.001) (**Figure S7b**). One possible explanation is that the high-affinity binders recruit p300 during transient dCas9 interactions at off-target sites, thereby activating transcription at these sites. TAZ2 binders reproducibly upregulated a small set of additional genes, including *SNCG*, *S100A1*, and *WFDC21P* **(Figure 4f, Figure S7b)**. GO analysis did not reveal a coherent functional signature, suggesting that these changes are unlikely to reflect activation of a specific transcriptional program. Because TAZ2 participates in autoinhibitory regulation of p300 acetyltransferase activity, high-affinity engagement of this domain could relieve autoinhibition and increase p300 activity more broadly. Thus, KIX- and TAZ1-targeting binders may be preferable for CRISPRa applications, given their limited transcriptome-wide perturbation and robust target activation.

Overexpression of conventional TADs can impair cellular fitness, potentially through nonspecific interactions or sequestration of transcriptional machinery^27^. We therefore asked whether the designed binders imposed a similar cellular burden. We compared lentiviral production from HEK293T cells packaging constructs encoding dCas9 alone, dCas9 fused to the designed binders, VPR, or the p300 catalytic core. Viral titers were quantified by transducing an equal number of recipient cells and measuring vector integration by qPCR. Across the five binders tested, titers were relatively consistent and approximately half that of dCas9 alone, but remained 3–6-fold higher than VPR and 10–20-fold higher than the p300 catalytic-core fusion **(Figure S7d)**. These results suggest that the designed binders impose substantially less cellular burden than conventional VPR or direct p300 fusion.

To confirm that the binders function across diverse genomic contexts, we tested selected binders at the promoters and enhancers of *OCT4* and *HBG*, along with several additional loci. Although activity varied by locus, the binders were generally comparable to or weaker than VPR at promoters, but consistently outperformed VPR at enhancers, highlighting the prominent role of p300 in enhancer-mediated gene activation **(Figure 4g-i, Figure S7e)**^28^. We reasoned that the stronger promoter activity of VPR could arise from its ability to engage multiple transcriptional coactivators, whereas the designed binders selectively recruit p300. To test whether additional coactivator recruitment could further enhance activity, we generated dual activators by fusing VP64 and the p300 binders to the N- and C-termini of dCas9, respectively **(Figure 4j)**. At the *OCT4* promoter, addition of VP64 increased activation by 3- to 24-fold across the three binders tested, exceeding both the corresponding binder alone and VPR **(Figure 4j)**. Together, these results show that the p300 binders function across diverse genomic loci, with particularly strong activity at enhancers, and that combining selective p300 recruitment with additional coactivator recruitment can further enhance transcriptional activation.

## Discussion

Transcriptional activation is associated with the recruitment of coactivators, but native activation domains make this process difficult to dissect quantitatively because their interactions are often weak and promiscuous. Here, we used structurally defined de novo binders to examine how coactivator recruitment is translated into transcriptional output. We found that cooperative recruitment of coactivators occurs at multiple levels, from distinct interfaces within a single domain to different domains of the same coactivator, and potentially across components of larger transcriptional complexes. The dual-interface TAZ2 binder provides a direct example in which two weak interactions with separate surfaces of one domain can produce strong activation. The KIX affinity series further demonstrated that weak KIX interactions can cooperatively recruit p300, and revealed a sharp, switch-like relationship between binding affinity and transcriptional output, with an apparent threshold around 500 nM. Weaker interactions were largely insufficient for activation, whereas stronger binding above this threshold provided little additional transcriptional benefit. Together, these results suggest that multivalent cooperativity may be a general feature of coactivator recruitment. Defining these principles more broadly should improve both our understanding of gene regulation and our ability to engineer it with greater precision.

Beyond their use as mechanistic probes, the designed binders provide a compact and modular platform for synthetic gene regulation. Potent yet compact p300 binders drove robust endogenous gene activation exceeding VPR while imposing much lower apparent cellular burden. More importantly, their defined specificity makes coactivator recruitment programmable: rather than relying on native activation domains that engage heterogeneous sets of coactivators, individual molecular functions can be recruited and combined explicitly. The enhanced activity observed when p300 recruitment was paired with additional coactivators shows how defined coactivator modules can be assembled to build stronger and more tunable transcriptional regulators. The compact size and defined specificity of these effectors could also facilitate multiplexed CRISPRa using Cas12a-based systems^29–31^, enabling systematic perturbation of regulatory programs and combinatorial dissection of gene regulatory networks. This modularity could extend beyond CRISPRa: fusing designed coactivator binders to native or de novo designed DNA binding domains^32,33^ could create hybrid or fully synthetic regulators that couple precise DNA recognition with chosen regulatory activities.

In summary, we have established a general strategy for designing and screening functional transcriptional regulators with defined molecular interactions. Although developed here for gene activation, the same framework could be extended to repression by targeting corepressors and silencing complexes, enabling systematic control over both activating and repressive pathways. Expanding the repertoire of designed regulators could ultimately allow gene expression to be tuned more continuously and combinatorially, rather than simply switched on or off. More broadly, programmable recruitment of defined regulatory activities may provide a path toward precise control of gene-expression programs, cell states, and cellular functions.

## Supplementary_Figures

**Figure S1.**
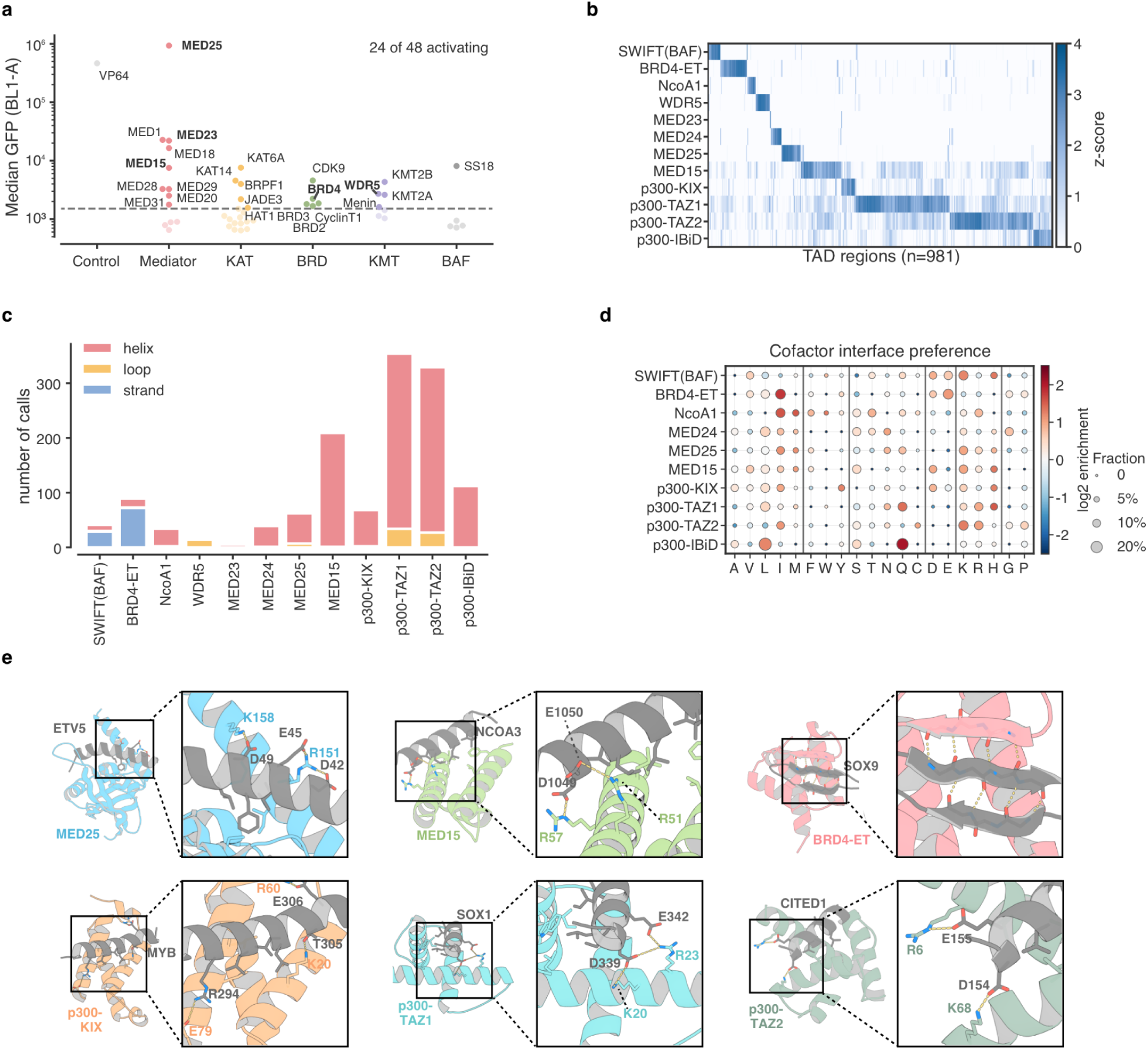
Functional and structural characterization of TAD–coactivator interactions. **a.** Transcriptional activity of representative coactivators considered for TAD–coactivator interaction mapping and binder design. Coactivators were grouped by functional class, including Mediator, lysine acetyltransferases (KAT), bromodomain-containing proteins (BRD), lysine methyltransferases (KMT), and BAF complex components. Representative coactivators were fused to the C terminus of a zinc-finger DNA-binding domain and co-transfected with a GFP reporter into HEK293T cells. Median GFP fluorescence was quantified by flow cytometry. The dashed line indicates the activation threshold; 24 of 48 coactivators tested showed detectable activation. **b.** Heatmap showing relative TAD interaction preferences across coactivator domains. AF3-predicted interactions were quantified by ipSAE and z-scored within each coactivator to emphasize relative preference across TADs. This analysis complements the overall TAD–coactivator interaction landscape shown in Figure 1d. **c.** Predicted secondary structures of TADs upon coactivator binding. Bars indicate the numbers of predicted helical, loop, and β-strand interaction modes for each coactivator domain. Most interactions were α-helical, whereas SWIFT and BRD4-ET interactions were enriched for β-strand conformations. **d.** Coactivator amino-acid preferences at predicted interfaces. Circle size indicates the fraction of interacting coactivators containing each residue at the interface, and color indicates log2 enrichment relative to proteome background. Most coactivators show enrichment in positively charged residues, complementing the sequence-preference analysis in Figure 1e. **e.** Representative AF3-predicted TAD–coactivator complexes illustrating distinct binding modes and polar interactions. Coactivators and TADs are shown in the indicated colors, with enlarged views highlighting representative interface residues and predicted electrostatic interactions.

**Figure S2.**
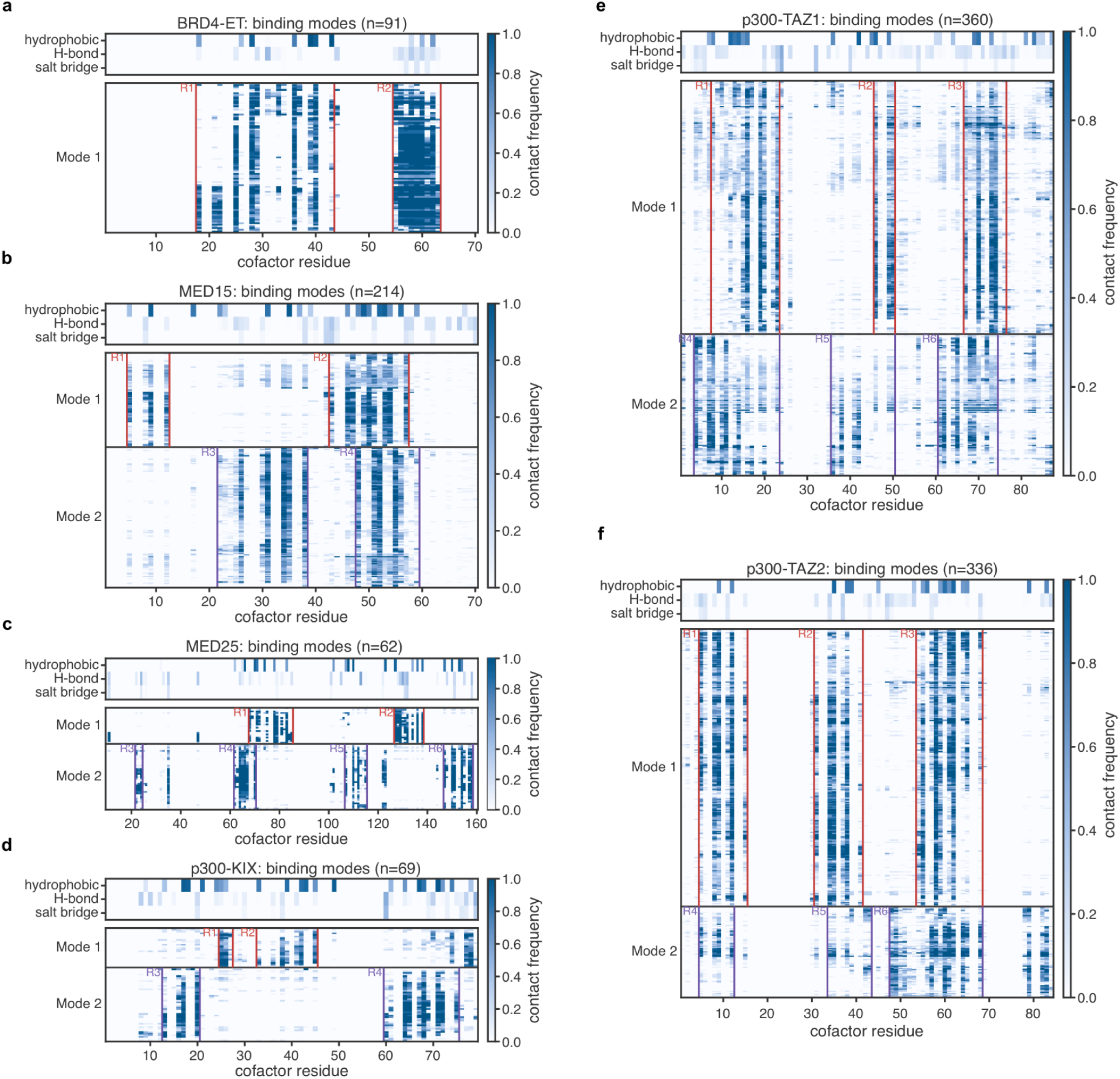
Predicted TAD-binding modes reveal multiple interaction surfaces on transcriptional coactivators. a–f, AF3-predicted TAD-binding modes for **BRD4-ET (a), MED15 (b), MED25 (c), p300-KIX (d), p300-TAZ1 (e), and p300-TAZ2 (f)**. Each row in the lower heatmaps represents a predicted TAD–coactivator interaction and each column a coactivator residue; color indicates contact frequency. Colored boxes mark recurrent contact regions associated with each mode. Upper heatmaps summarize the frequencies of hydrophobic contacts, hydrogen bonds, and salt bridges at each coactivator residue. These predicted binding modes were used to select target interfaces and key residues for RFdiffusion-based binder design. The presence of multiple recurrent surfaces motivated subsequent tests of whether individual coactivator interfaces act independently or cooperatively in **Figure 2f–i**.

**Figure S3.**
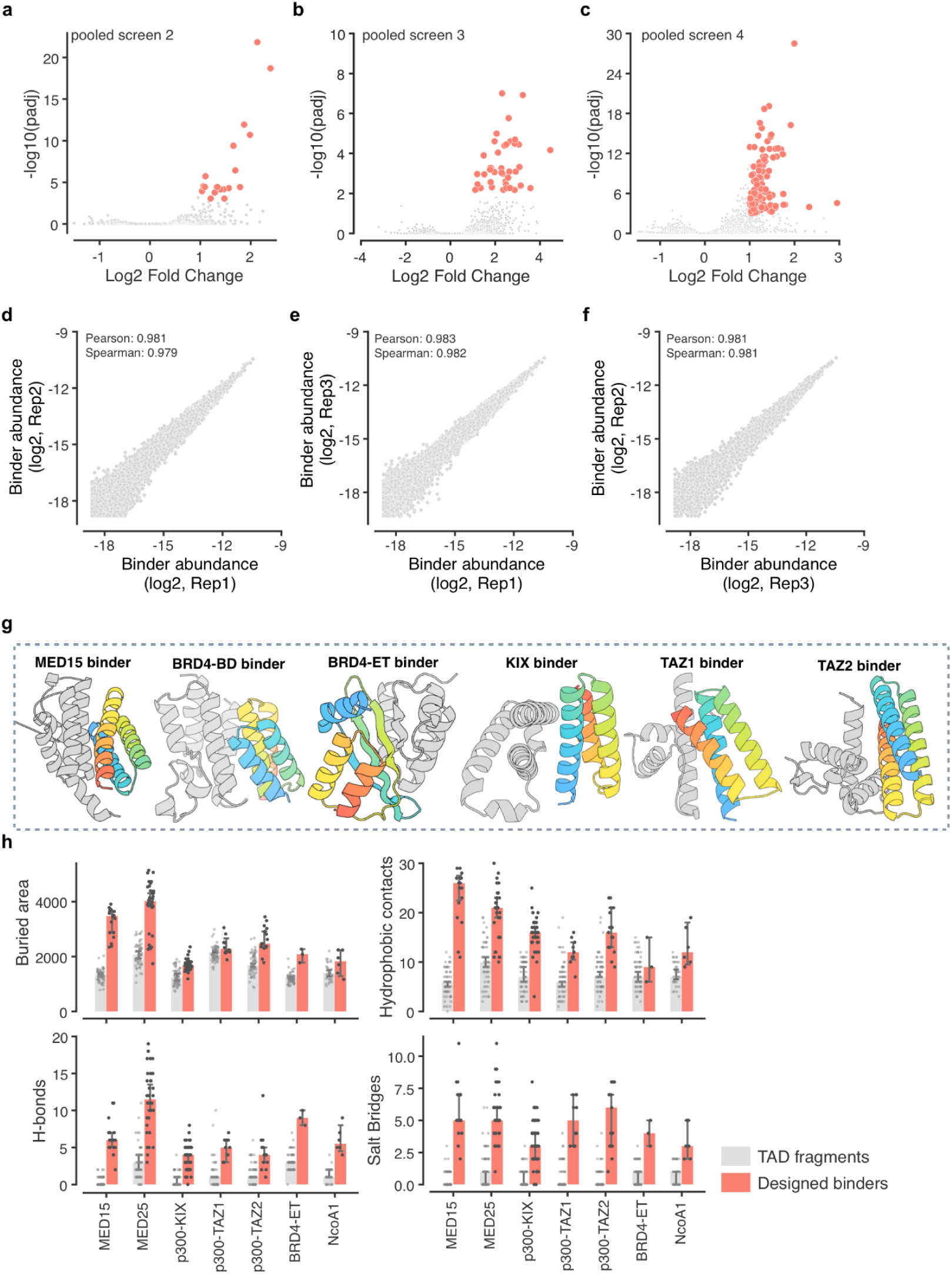
Pooled screening and characterization of designed binders. a–c, Volcano plots show binder enrichment after functional selection. Red points indicate significant and substantial enrichment (Wald-test with Benjamini-Hochberg correction P <0.001 for fold-difference >2). **d–f,** Reproducibility of pooled screens across independent integration replicates. Log-scaled abundances of individual binders are highly correlated across replicates. **g,** Representative AF3-predicted structures of experimentally identified coactivator binders. Target coactivators are shown in gray and designed binders in color. **h,** Comparison of predicted interfaces formed by native TAD fragments and designed binders across the indicated coactivators. Interface properties include buried surface area, hydrophobic contacts, hydrogen bonds, and salt bridges. Designed binders generally form larger interfaces with more extensive hydrophobic and polar interactions than native TAD fragments.

**Figure S4.**
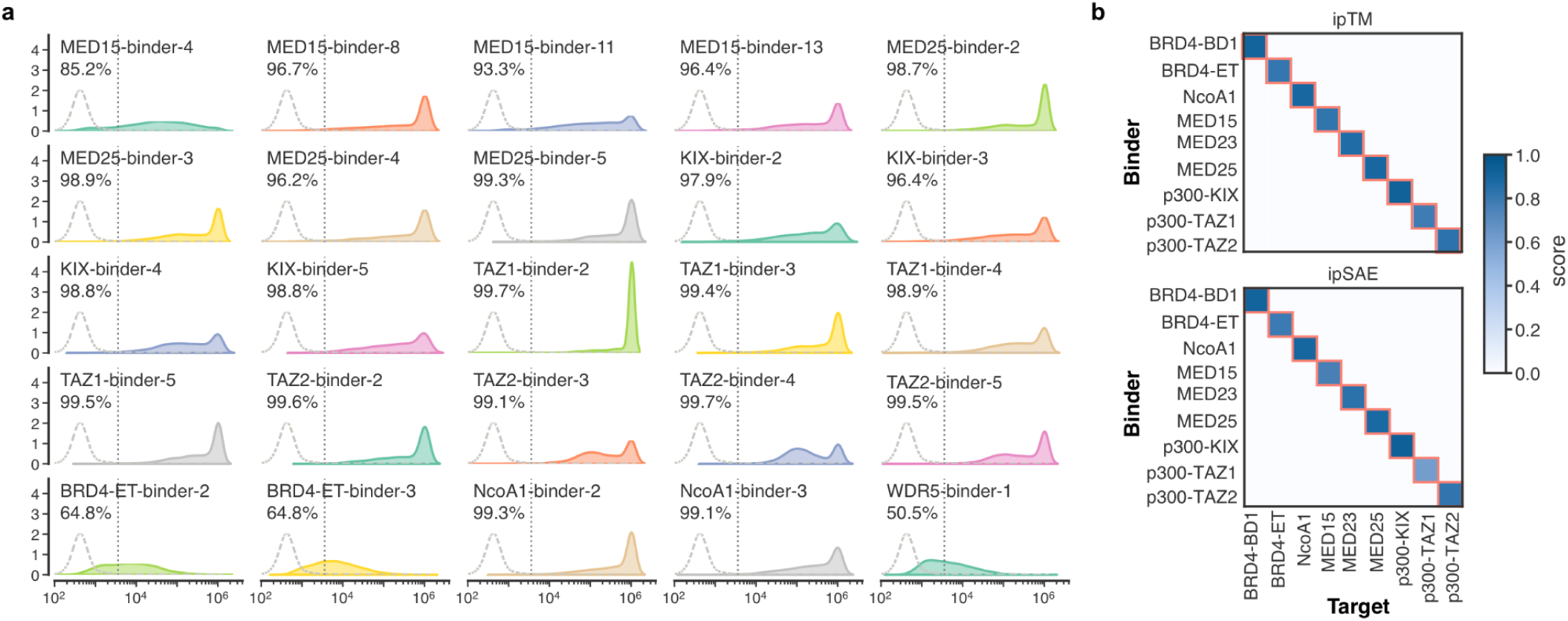
Functional validation and predicted specificity of designed coactivator binders. **a.** Histogram of GFP fluorescence intensity for the indicated binder activators. Extended panel of **Figure 2c**. **b.** AF3-predicted specificity of designed binders. Binder–coactivator pairs with predicted contact probability <0.5 were masked. Interaction confidence is shown by ipTM and ipSAE scores, providing complementary specificity metrics to the contact probability shown in **Figure 2e**.

**Figure S5.**
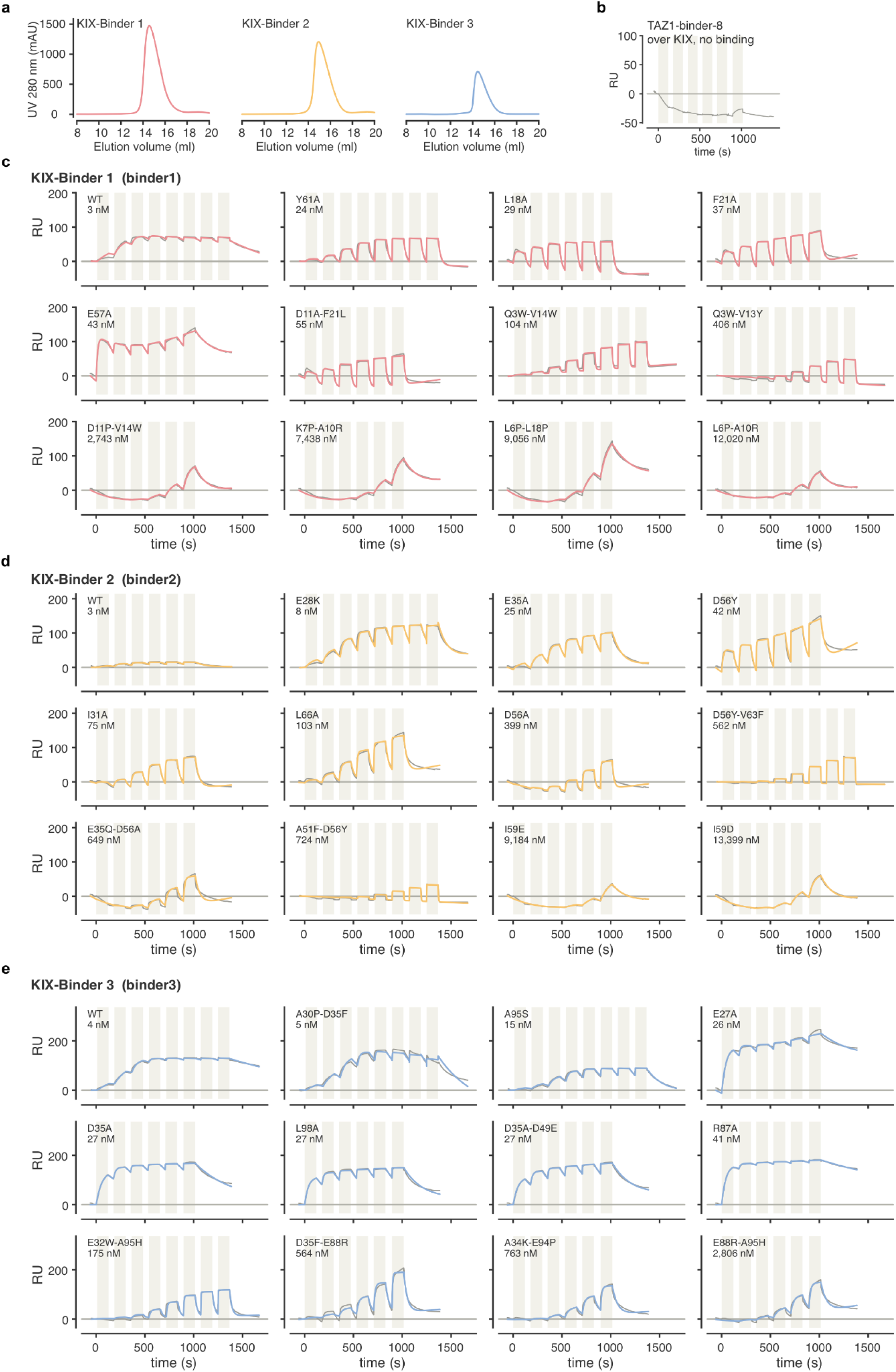
Biochemical characterization of affinity-tuned KIX binders. a, Size-exclusion chromatography profiles of purified KIX-Binders 1–3. Each binder eluted as a predominant monomeric peak, consistent with soluble, monodisperse proteins. **b,** SPR specificity control using a TAZ1 binder injected over immobilized KIX, showing no detectable KIX binding. **c–e,** Complete SPR sensorgrams for the parental proteins and affinity-tuned variants of **KIX-Binder 1 (c), KIX-Binder 2 (d), and KIX-Binder 3 (e)**. Measured K_D_ values are indicated for each variant.

**Figure S6.**
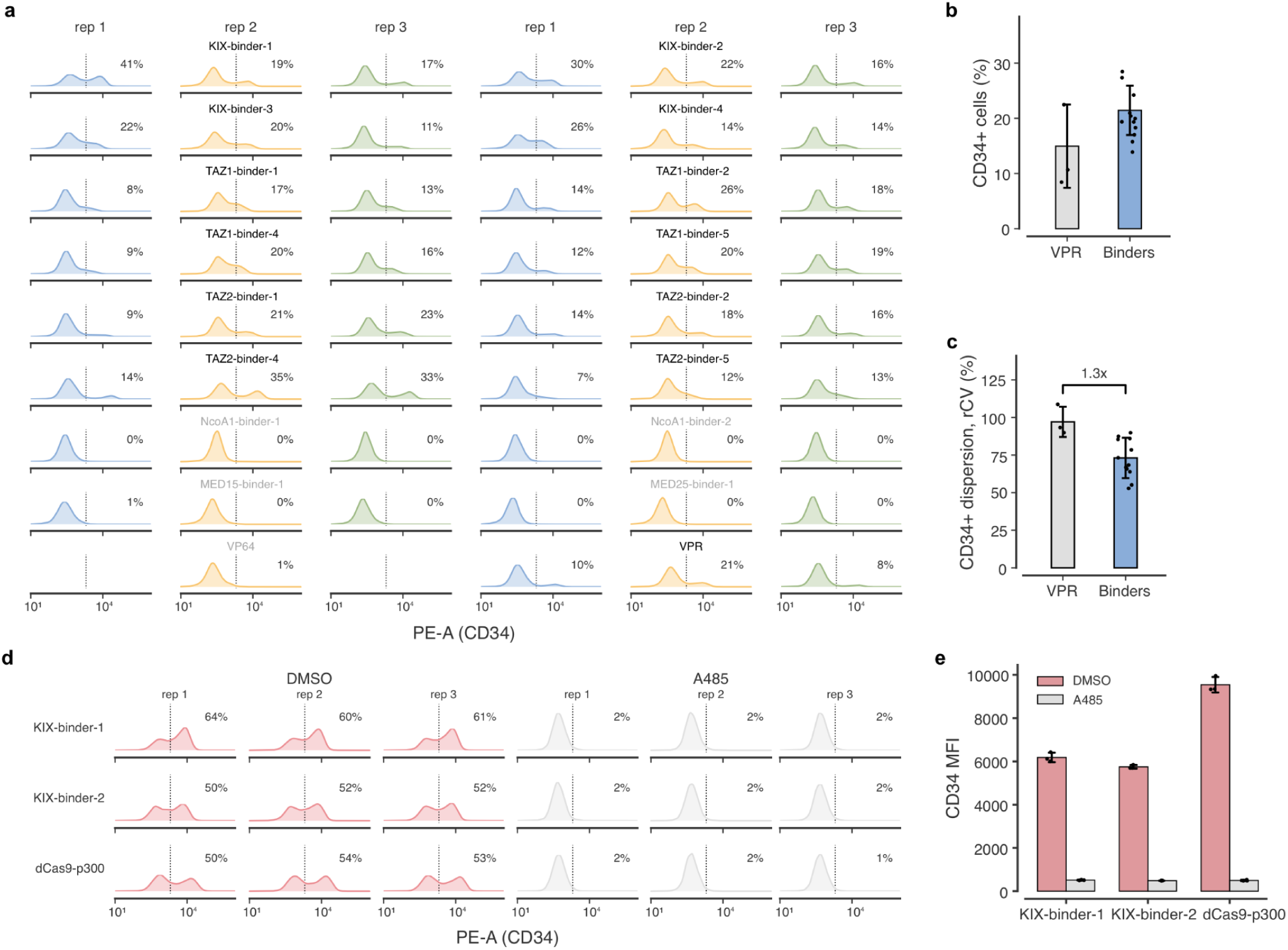
CRISPRa with designed binders. **a.** Endogenous CD34 activation by dCas9-binder fusion in triplicates. Related to **Figure 4b**. **b.** Fraction of CD34-positive cells following activation by VPR or the designed p300 binders. Points represent individual measurements. **c.** Dispersion of CD34 expression within the activated population, quantified by robust coefficient of variation (rCV). Designed binders produced lower expression variability than VPR. **d.** Requirement for p300 catalytic activity. Flow-cytometry distributions of CD34 expression following activation by KIX-Binder 1, KIX-Binder 2, or dCas9–p300 in cells treated with DMSO or the p300 catalytic inhibitor A-485. Three independent replicates are shown. **e,** Quantification of CD34 mean fluorescence intensity (MFI) for the conditions shown in **d**. A-485 strongly suppressed activation by both KIX binders and direct p300 recruitment, indicating that binder-mediated CD34 activation requires p300 catalytic activity.

**Figure S7.**
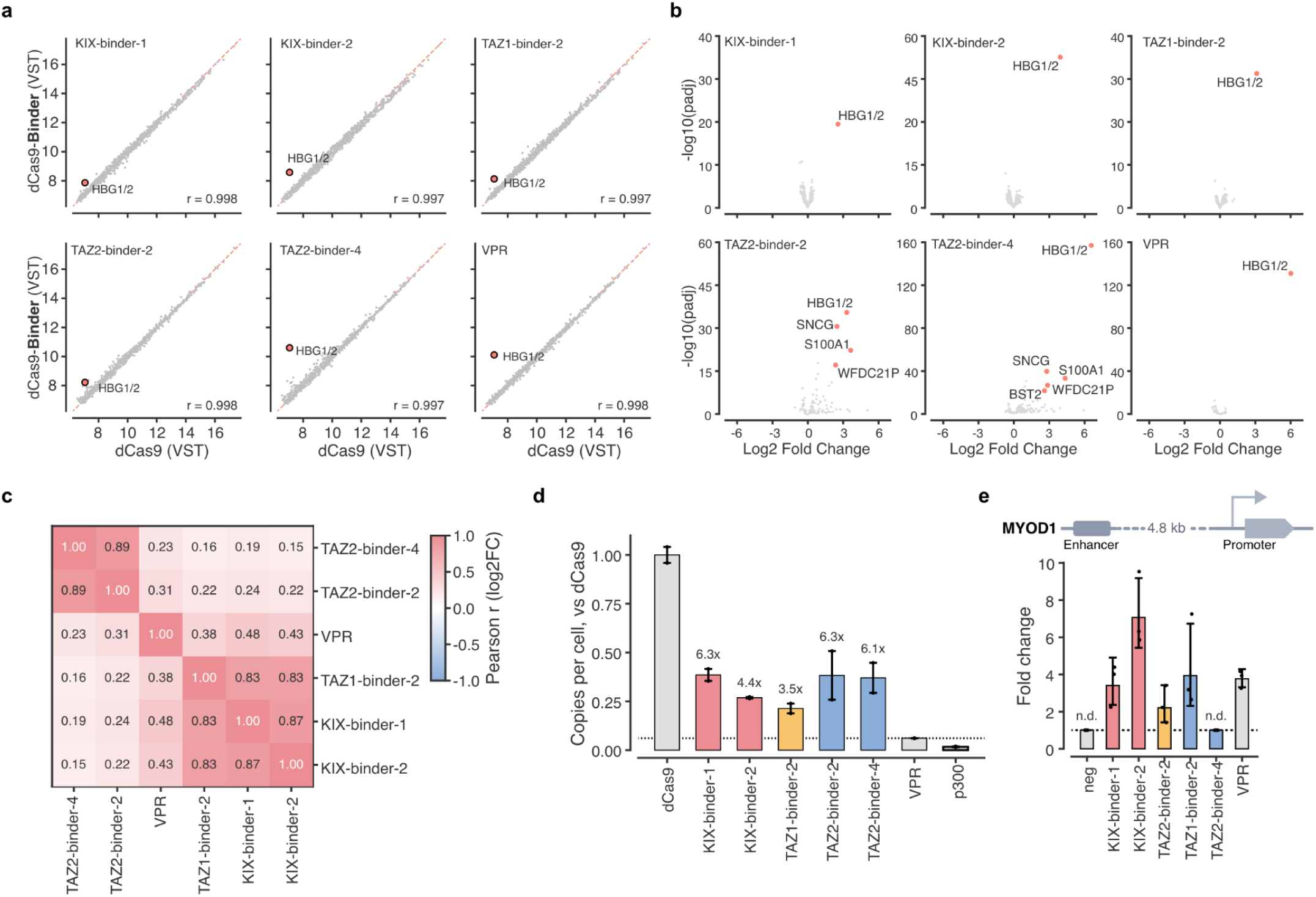
Characterizing specificity and cell burdens of designed binders. a, Comparison of transcriptome-wide gene expression between dCas9 alone and the indicated dCas9–activator fusions following targeting of the HBG locus. Variance-stabilized transformed (VST) expression values are shown for all genes, with HBG1/2 highlighted. Pearson correlation coefficients are indicated. **b,** Volcano plots of differentially expressed genes in cells expressing dCas9-binders with sgRNA targeting the HBG locus. HBG1/2 was strongly activated by all indicated p300 binders and VPR. Complementary to **Figure 4 e,f. c**, Similarity of transcriptome-wide responses induced by the different activators. Heatmap shows pairwise Pearson correlations between gene-level log2 fold changes. KIX- and TAZ1-targeting binders show highly similar transcriptional profiles, whereas TAZ2 binders form a distinct group. **d.** Relative cellular burden of dCas9 activators assessed by lentiviral production. Viral particles produced from cells expressing dCas9 alone, the indicated p300 binders, VPR, or a p300 fusion were used to transduce recipient cells, and vector integration was quantified as copies per cell. Designed binders supported substantially higher viral production than VPR or direct p300 fusion. **e.** Activation of MYOD1 by targeting its enhancer located approximately 4.8 kb upstream of the promoter.

## Contributions

W.C. and D.B. conceptualized the project. W.C. designed the binders with assistance from B.C., M.B., K.W., and C.G. W.C., R.J., I.B.H., and D.B. designed the experiments. W.C. and R.J. performed the experiments with assistance from X.Q., M.B., L.T., T.T., Z.S., N.R., and X.L. W.C. analyzed the data with input from I.B.H. and M.B. W.C. and D.B. wrote the manuscript with input from all authors. D.B. supervised the project.

## Acknowledgements

We thank members of the Baker Lab, particularly Ian Humphreys, Nikita Hanikel, Han Raut Altae-Tran, Bo Yang, Thomas Schlichthärle, Basile Wicky, Shunzhi Wang, as well as Bo Gu from M. Elowitz lab at Caltech for helpful discussions. We thank K. VanWormer, and L. Goldschmidt for general technical support. We thank D. Liu’s lab for sharing prime editing plasmids. This work was supported by a grant from The Audacious Project at the Institute for Protein Design (PG117878, PG117866 to D.B.), National Institute of General Medical Sciences (R35GM143532 to I.B.H.) and Welch Foundation (C-2240-20250403 to I.B.H.). W.C. is a Fellow of the Damon Runyon Cancer Research Foundation (DRG-2507-23). D.B. is an Investigator of the Howard Hughes Medical Institute.

## Data availability statement

Raw sequencing data and associated processed data have been deposited to Gene Expression Omnibus (GEO) with accession numbers GSE351295 and GSE351296. Custom scripts and Jupyter notebooks related to this paper are available at GitHub: https://github.com/chenomics/synTAD_design.

## Disclosures

The authors used AI tools to assist with data analysis and improve the clarity and conciseness of the manuscript. All AI-assisted analyses and text were reviewed and edited by the authors.

## Supplementary Table

**Supplementary Table 1 | List of 726 TADs used for AF3 predictions and 981 TAD regions with their interaction partners**

**Supplementary Table 2 | List of coactivator domains used for AF3 predictions and binder design**

**Supplementary Table 3 | Sequences of primers and other oligos**

## Materials and Methods

### AF3 prediction on the TAD-coactivator interactions

TADs were curated from published screens and reviews^3,4,14^ and tiled into 50-aa windows with a 10-aa step, yielding 6,520 TAD windows. Interactions were predicted against 13 coactivator domains, generating 84,760 TAD–coactivator pairs. TAD windows were provided to AlphaFold 3 (AF3) as single sequences without MSAs or structural templates. For each coactivator domain, an MSA was generated using HHblits^34^ against UniRef30_2022_02 with four iterations and an E-value cutoff of 1e-10, then filtered using hhfilter at 98% maximum pairwise identity and 50% minimum coverage. The resulting MSAs contained approximately 330–52,000 sequences, with most containing 600–2,700 sequences. Each coactivator chain was additionally provided with a structural template derived from the corresponding AlphaFold model, with query and template residues explicitly mapped. No paired MSAs were used. For p300 TAZ1 and TAZ2, three Zn^2+^ ions were included as ligands. All complexes were predicted using a local installation of AF3 with the data pipeline disabled and precomputed MSAs and templates, using one model seed and five diffusion samples per seed. TAD sequences are listed in **Supplementary Table 1**.

### Designing coactivator specific binders

Binder backbones were generated using RFdiffusion3. For each design, the target domain was supplied as a single structural chain, with its sequence and residue indices held fixed. A second chain was generated to form a two-chain binder–target complex. Binder lengths were sampled from 60-100 residues. Because RFdiffusion3 samples a single binder length per batch, each design campaign consisted of 100 independent batches of 10 designs, with a new length sampled for each batch. In total, 1,000 backbones were generated per target.

Binder sequences were designed using ProteinMPNN, with the target sequence held fixed and all binder residues redesigned. For each backbone, 10 sequences were sampled at a temperature of 0.1. To reduce alanine enrichment at solvent-exposed positions, we applied an alanine bias of −1.0 to residues identified as exposed using a geometry-based, direction-weighted neighbor count. We also applied a methionine bias of −8.0 at the first binder position to discourage N-terminal methionine incorporation.

Designed sequences were filtered based on predicted binding energy calculated using PyRosetta. Each sequence was threaded onto its backbone, and the binder and neighboring target residues within 10 Å were repacked using the beta_nov16 energy function. Binding energy was calculated as the difference in soft-repulsive energy between the repacked complex and the separated chains, without additional repacking after separation. Designs with predicted interface ΔΔG < −24 were selected for further evaluation.

Selected binder–target complexes were predicted using AlphaFold3 (AF3). The target structure was provided as a template while the binder was modeled without a template. No MSA was used in the prediction. Each complex was predicted using 10 recycles and five diffusion samples. Designs were retained if they met the following criteria: interface predicted TM-score (ipTM) > 0.85, minimum interface predicted aligned error (PAE) < 1 or 1.5 Å, mean interface PAE < 7 Å, and interface Cα RMSD < 15 Å between the predicted and designed structures.

Designs passing the AF3 filters were further diversified through partial diffusion using RFdiffusion3. To avoid redundant sampling, designs originating from the same backbone were grouped, and one representative was selected for each backbone. The AF3-predicted complexes were then subjected to partial diffusion with a maximum noise level of 10 Å, allowing local structural variation while largely preserving the overall fold and binding orientation. During partial diffusion, the target structure was held fixed while the binder backbone was regenerated. Each resulting backbone underwent independent sequence design, interface energy filtering, and AF3 prediction using the same procedures and selection criteria described above.

### Molecular cloning

Primers and oligonucleotides used in this study are listed in **Supplementary Table 3**. Unless otherwise indicated, PCR products and digested plasmid DNA were purified using AMPure XP beads (Beckman Coulter) at bead-to-sample volume ratios of 1.2x and 0.6x, respectively, according to the manufacturer’s instructions. Plasmid DNA used for mammalian-cell experiments was prepared using the ZymoPURE II Plasmid Kit (Zymo Research).

Binder library cloning: Pooled DNA oligonucleotides encoding the designed binder libraries were synthesized by Twist Bioscience. Libraries were amplified using Q5 Hot Start High-Fidelity DNA Polymerase (NEB) with the following cycling conditions: 98 °C for 3 min; 22 cycles of 98 °C for 20 s, 65 °C for 15 s, and 72 °C for 60 s. PCR products were purified, digested with BsmBI, and purified again. The destination plasmid (P102) was separately digested with BsmBI and purified. Digested binder inserts and destination vector were ligated using T4 Ligase (NEB) at a 1:6 vector to insert molar ratio. To maximize library complexity, each ligation was electroporated into NEB 10-beta Electrocompetent *E. coli* (C3020) in duplicate. Following recovery, transformed cells were combined and cultured in 50 mL medium at 30 °C overnight. Plasmid libraries were subsequently purified for mammalian-cell transfection.

Individual binder validation cloning: For individual validation, DNA fragments encoding the designed binders flanked with the BsaI sites and assembly overhangs were synthesized by Twist Bioscience. Constructs were assembled into the corresponding destination vectors (P171) by one-step Golden Gate assembly using BsaI, T4 DNA Ligase Buffer, and T4 DNA Ligase (New England Biolabs) in a total reaction volume of 15 µL. Insert and destination vector were used at a 1:1 molar ratio. Assembly reactions were incubated at 37 °C for 60 min followed by 60 °C for 5 min. Assembly products were transformed into NEB Stable Competent *E. coli* (C3040) according to the manufacturer’s protocol and plated on selective agar. Individual colonies were picked for plasmid preparation and downstream validation.

### Cell culture, transient transfections, nucleofection and piggyBac integrations

HEK293T (CRL-11268) and K562 cells (CCL-243) were purchased from ATCC. HEK293T cells were cultured in DMEM, high glucose (GIBCO). K562 cells were cultured in RPMI 1640 medium (GIBCO). All media were supplemented with 10% Fetal Bovine Serum (Hyclone) and 1% penicillin-streptomycin (GIBCO). Cells were grown with 5% CO_2_ at 37°C. Transfection of HEK293T, K562 cells was performed using Lipofectamine 3000 (ThermoFisher, L3000015), Lonza 4D-Nucleofector, respectively, following the manufacturer’s protocol.

Generation of the recorder screening cell line: The recorder screening cell line was generated by stable integration of a TRE–ENGRAM recorder and a genomic DNA tape containing a frameshifted puromycin-resistance open reading frame into HEK293T cells using the piggyBac transposon system. 1 x 10^6^ Cells were seeded in 6-well plates and transfected with 1.6 µg of the piggyBac cargo plasmid and 400 ng of Super piggyBac transposase expression plasmid (System Biosciences), for a total of 2 µg plasmid DNA per well. Following integration and selection, single cells were deposited into 96-well plates using a cell sorter and expanded clonally. Individual clones were screened for recorder activity, and a monoclonal line with robust and reproducible recording efficiency was selected for subsequent pooled screening experiments.

Integration of binder libraries for pooled screening: For each replicate of the pooled screen, 3 x 10^6^ cells were transfected with 5 µg binder-library plasmid and 500 ng Super piggyBac transposase expression plasmid. Three independent integrations were performed for each library and maintained separately in three culture flasks. Following transfection, cells were expanded for 7 days before initiation of the screening assay to allow dilution of residual episomal plasmid DNA.

Transient transfection for individual binder validation: For individual validation assays, 5 x 10^4^ cells were seeded per well in 96-well plates 24 h before transfection. Cells were co-transfected with 50 ng plasmid encoding the binder construct and 100 ng reporter plasmid. Cells were harvested 24 h after transfection and analyzed by flow cytometry.

Transient transfection for gene activation: For CRISPRa experiments, 1.5 x 10^5^ cells were seeded per well in 24-well plates 24 h before transfection. Cells were co-transfected with 187.5 ng of the indicated effector expression plasmid and 125 ng of sgRNA expression plasmid or sgRNA plasmid pool. Cells were collected 72 h after transfection for flow cytometry, RT-qPCR, or RNA sequencing.

### Binder sequence retrieval and quantification

To quantify binder abundance before and after pooled screening, cells were collected immediately before and after puromycin selection. Approximately 1 x 10^7^ cells before selection and 3 x 10^6^ cells after selection were harvested for each replicate, and genomic DNA was purified.

Integrated binder sequences were recovered by two-step PCR using KAPA2G Robust HotStart ReadyMix. To maintain sufficient library representation for accurate quantification of binder abundance, 24 parallel first-round PCR reactions were performed for each pre-selection sample and 8 reactions for each post-selection sample. In the first PCR, the binder-encoding region was amplified from genomic DNA using 0.5 µM forward and reverse primers in a 50-µL reaction. PCR was performed at 95 °C for 3 min, followed by 22 cycles of 98 °C for 20 s, 65 °C for 15 s, and 72 °C for 40 s. PCR products were size-selected using sequential 0.5× and 1× AMPure XP bead purification to remove genomic DNA and small DNA fragments, respectively. For the second PCR, 1 ng of purified first-round PCR product was amplified using 0.2 µM primers containing Illumina flow-cell adapters and sample indices in a 25-µL reaction. PCR was performed at 95 °C for 3 min, followed by 5 cycles of 98 °C for 20 s, 65 °C for 15 s, and 72 °C for 40 s. Indexed PCR products were pooled, purified using AMPure XP beads, and sequenced on an Illumina platform. Binder abundance in each sample was determined from sequencing read counts and used to quantify enrichment following puromycin selection.

### Sequencing data analysis and binder enrichment

Sequencing reads from pooled binder screens were aligned to the reference binder sequence library using BWA (version 0.7.1)^35^ with default settings. Aligned reads were processed using custom scripts to count the number of reads corresponding to each binder in each sample.

Raw binder counts from pre- and post-puromycin-selection samples were used as input for DESeq2^36,37^. Differential enrichment was assessed using the Wald test, and P values were adjusted for multiple testing using the Benjamini–Hochberg procedure. Binders were classified as enriched if they showed greater than twofold enrichment after puromycin selection and an adjusted P value <0.001.

### Flow cytometry

For individual binder validation, cells were collected 24 h after transfection. Cells were detached by trypsinization, pelleted, and resuspended in FACS buffer consisting of 1% FBS in PBS. Samples were analyzed on an Attune flow cytometer.

For CRISPRa experiments, cells were collected 72 h after transfection. Cells were detached by trypsinization, pelleted, and resuspended in FACS buffer consisting of 1% FBS in PBS. For CD34 expression analysis, cells were washed twice with 200 µL ice-cold FACS buffer and resuspended in 25 µL PE-conjugated anti-CD34 antibody solution (Invitrogen, clone QBEND/10, MA1-10205) diluted 1:20 in FACS buffer. Cells were incubated for 30 min at 4 °C protected from light, washed twice with 200 µL FACS buffer, and resuspended in 200 µL FACS buffer for acquisition on a Sony SA3800 spectral analyzer.

Flow-cytometry data were analyzed using the custom floway package. Cell debris was excluded based on FSC-A and SSC-A, followed by singlet selection using FSC-A and FSC-H. GFP-positive gates were established with reporter plasmid only samples, and fluorescence intensity was reported as median of GFP intensities. CD34-positive gates were established using unstained controls. CD34 positivity was reported as the percentage of singlets, and fluorescence intensity was reported as the geometric mean PE fluorescence of CD34-positive singlets.

### Reverse-transcription quantitative PCR

Cells were collected 72 h after transfection or nucleofection, unless otherwise indicated, and total RNA was isolated using the RNeasy Plus Mini Kit (Qiagen, #74136). RNA concentrations were measured using a NanoDrop spectrophotometer, and 1 µg of RNA was used for cDNA synthesis with the iScript Advanced cDNA Synthesis Kit for RT-qPCR (Bio-Rad, #1725038). cDNA was diluted 10-fold with nuclease-free water. Each 20-µL qPCR reaction contained 10 µL of SsoAdvanced Universal SYBR Green Supermix (Bio-Rad, #1725275), 9 µL of diluted cDNA, and 1 µL of gene-specific forward and reverse primer mix. Reactions were performed using a CFX96 Real-Time PCR Detection System with a C1000 thermal cycler (Bio-Rad). Target-gene expression was normalized to GAPDH, and relative expression was calculated using the 2^−ΔΔCt method relative to the control. Primer sequences are provided in **Supplementary Table 3**.

### Lentivirus production

Lentiviral preparations were generated in HEK293T cells cultured in 10-cm dishes by co-transfection of the indicated transfer plasmid with psPAX2 and pMD2.G. Transfections used linear polyethylenimine (PEI 25K; Kyfora Bio, #23966) and Opti-MEM. Following transfection, cultures were maintained in complete DMEM supplemented with sodium butyrate. Virus-containing supernatants from successive collections were pooled, clarified, and filtered using PES syringe filters (Puradisc PES 25AS; VWR, #28137-938). Preparations were concentrated using Lenti-X Concentrator (Takara, #631232).

### Lentivirus Functional Titration

HEK293T cells were plated in a 24-well plate at 150,000 cells/well directly with concentrated lentivirus at 1% v/v in Opti-MEM. 72 hours after transduction, DNA was isolated using the DNeasy Blood & Tissue Kit (QIAGEN, #69506). Vector-associated WPRE DNA and the human albumin reference locus (ALB) were quantified by qPCR using SsoAdvanced Universal SYBR Green Supermix (Bio-Rad, #1725275) on the CFX96 system. Copy numbers were estimated by reference to a WPRE/ALB plasmid standard curve. Vector copies per cell were estimated using the WPRE copy number, normalized to the ALB copy number. Functional titers were calculated from the normalized copy estimates, recipient-cell input, and volume of preparation tested. Primer sequences are provided in **Supplementary Table 3**.

### Bulk RNA-sequencing

HEK293T Cells were collected 72 h after transfection in 50 µL of 1X DNA/RNA Shield (Zymo Research, R1100-50) per sample, and three biological replicates per condition were submitted to Plasmidsaurus for RNA sequencing. Gene-level fractional counts were supplied by the sequencing provider. Each of the experimental conditions was compared separately with the dCas9-only controls, yielding 6 contrasts. P-values were adjusted using the Benjamini–Hochberg procedure separately within each contrast. Differential expression analysis was performed using DESeq2^36,37^. Differentially expressed genes were called with thresholds of adjusted P <0.001 for a change > 2; positive fold changes indicate higher expression in the experimental condition.

### Recombinant protein expression and purification

Genes encoding the designed binders and binder variants were synthesized by Twist Bioscience, cloned into bacterial protein-expression vectors, and transformed into *E. coli* C2530H. Transformed cells were plated on selective agar, and individual colonies were picked and cultured overnight. Overnight cultures were diluted 1:100 into auto-induction medium and grown at 30 °C for 24 h.

Cells were harvested by centrifugation and resuspended in lysis buffer containing 50 mM Tris-HCl, pH 8.0, and 300 mM NaCl, supplemented with protease inhibitor cocktail immediately before use according to the manufacturer’s instructions. Cells were lysed by sonication, and insoluble material was removed by centrifugation at 14,000g for 30mins. His-tagged proteins were purified by immobilized metal-affinity chromatography. After binding, the resin was washed with buffer containing 50 mM Tris-HCl, pH 8.0, 300 mM NaCl, and 30 mM imidazole. Bound proteins were eluted with 50 mM Tris-HCl, pH 8.0, 300 mM NaCl, and 300 mM imidazole. Proteins were further purified by size-exclusion chromatography using either an HPLC or ÄKTA chromatography system equilibrated in 1× HBS-EP+ buffer. Fractions corresponding to the main monomeric peak were pooled. Purified protein concentrations were determined by NanoDrop and used for subsequent biophysical characterization.

### Surface plasmon resonance

Binding affinities of the purified binders were measured by surface plasmon resonance using 1× HBS-EP+ as the running buffer. Biotinylated p300 KIX protein was immobilized on a streptavidin-coated sensor surface, and purified binder proteins were used as analytes. For each binder, 6–8 concentrations were prepared as a threefold serial dilution in HBS-EP+ buffer and injected over the KIX-containing sensor surface. Concentration ranges were adjusted as needed to capture the binding transition for variants spanning different affinity regimes. SPR sensorgrams were reference-subtracted and independently fitted to a 1:1 single-cycle kinetic binding model. Five parameters were fitted for each sensorgram: the association rate constant k_a_, dissociation rate constant k_d_, maximum binding response R_max_, linear baseline drift, and exponential baseline decay amplitude. Fit residuals were evaluated at 0.5-s intervals across the injection series, excluding ±4 s around each injection start and stop. Equilibrium dissociation constants were calculated from the fitted kinetic parameters as K_D_=k_d_/k_a_.

## Notes

### Competing Interest Statement

The authors have declared no competing interest.

